# An AI-designed adenine base editor

**DOI:** 10.1101/2024.04.28.591233

**Authors:** Ye Yuan, Yang Chen, Rui Liu, Gula Que, Yina Yuan, Guipeng Li

## Abstract

Adenine base editors (ABEs) allow the efficient programmable conversion of adenine to guanine without causing DNA double strand breaks. Previous ABEs were generated by multiple rounds of directed evolution or derived by rational design based on the evolved ones. Although powerful, these methods search the local space for ABEs optimizations. Artificial intelligence (AI) based methods have the ability to efficiently explore much larger protein space for protein design. But currently there is no AI-designed ABE with wet experimental validation. Here, we demonstrate the first successful AI-designed ABE, which is named ABE10. ABE10 includes an AI-designed adenine deaminase enzyme fused with SpCas9n. The sequence identity between AI-designed enzyme and other publicly accessible variants is as low as 65.3%. ABE10 shows improved editing efficiency compared to current state-of-the-art ABE8 at multiple human genome sites tested. ABE10 also shows low off-target editing rate and reduced cytosine bystander effect. Our work demonstrates new direction for optimization of gene editing tools.

## Introduction

Adenine base editors (ABEs) are powerful tools in the realm of genome editing, specifically designed to make precise changes in the DNA sequence by converting adenine (A) to guanine (G) without inducing double-strand breaks^1^. ABEs offer a promising alternative to traditional CRISPR-Cas9 systems, which rely on the introduction of DNA double-strand breaks followed by error-prone repair mechanisms. ABEs comprise a catalytically impaired version of the CRISPR-Cas9 protein fused to an adenine deaminase enzyme TadA. The Cas9 protein and sgRNA compl guides the ABE to a specific target site within the genome, while the adenine deaminase enzyme catalyzes the conversion of adenine to inosine, which is read as guanine by DNA replication machinery, resulting in an A-to-G base conversion. One of the primary advantages of ABEs is their ability to induce precise single-base changes without causing large-scale disruptions to the DNA structure^2^. This precision makes them particularly valuable for correcting point mutations associated with genetic diseases or for engineering specific traits in organisms. ABE that disrupt PCSK9 are being evaluated in the clinic to treat atherosclerotic cardiovascular disease.

The deoxyadenosine deaminases in ABEs are derived from the Escherichia coli transfer RNA-specific adenosine deaminase (TadA)^3^. Through multiple rounds of directed evolution, several TadA variants were obtained, which correspond to the key component of the state-of-the-art ABEs, such as ABE7.10^1^, ABEmax^4^, ABE8.20^5^, ABE8e^6^ and ABE8r^7^. Based on ABE8e, two amino acids were mutated to obtain a new variant, ABE9^8^, which shows a narrow editing window with minimal bystander effect.

The applications of deep learning in protein design are increasing in recent years. Several deep learning methods have been developed for either fixed-backbone protein design or de novo protein design, such as ProteinMPNN^9^, RFdiffusion^10^, PiFold^11^, DIProT^12^ and so on. Experimentally characterized metal-binding protein and protein binders were designed with RFdiffusion^10^. PROTLGN was used to score the mutants of protein of interest and the top mutants were experimentally tested the function, such as fluorescence Intensity, binding affinity of VHH antibody, and cleavage activity of KmAgo^13^. The Graphormer-based Protein Design (GPD) model was employed to design CalB hydrolase with higher catalytic activity compared to the wild-type CalB^14^. There is a large potential for deep learning to accelerate the design process and reduce the number of variants that require experimental testing for protein function.

However, currently there is no AI-designed ABE reported with wet experimental validation. To achieve this goal, we developed a protein sequence design framework to learn the complex relationship between protein structure and amino acid sequence. Then we applied this framework to design the adenine deaminase enzyme, TadA, which is one key component of ABE. Ten sequences were selected and synthesized after codon optimization. The synthesized DNA are cloned to replace the tadA-8.20 DNA sequence in ABE8.20 plasmid, resulting in 10 ABE variants, which were tested in human cells. Two out of the 10 variants have base editing activity. One of them, named ABE10, shows significantly higher base editing efficiency at the test target site compared with ABE8.20. ABE10 also show higher editing efficiency at multiple sites with reduced Off-target and bystander effect.

## Results

### Protein design framework generates diverse adenine deaminase enzymes

The protein design framework is shown in Figure 1a, where protein design model and protein language model are integrated with a fusion module. The protein design model was trained as a fixed-backbone protein sequence design task, where the inputs are protein structures and outputs are protein amino acid sequences expected to fold into the input structures. The protein language model was trained as a masked language model task, which predicts the amino acid types of the random masked tokens in the input sequences. A fusion module that takes the output head of two models as input and predicts the distribution of amino acid types for the protein of interest.

**Figure 1.**
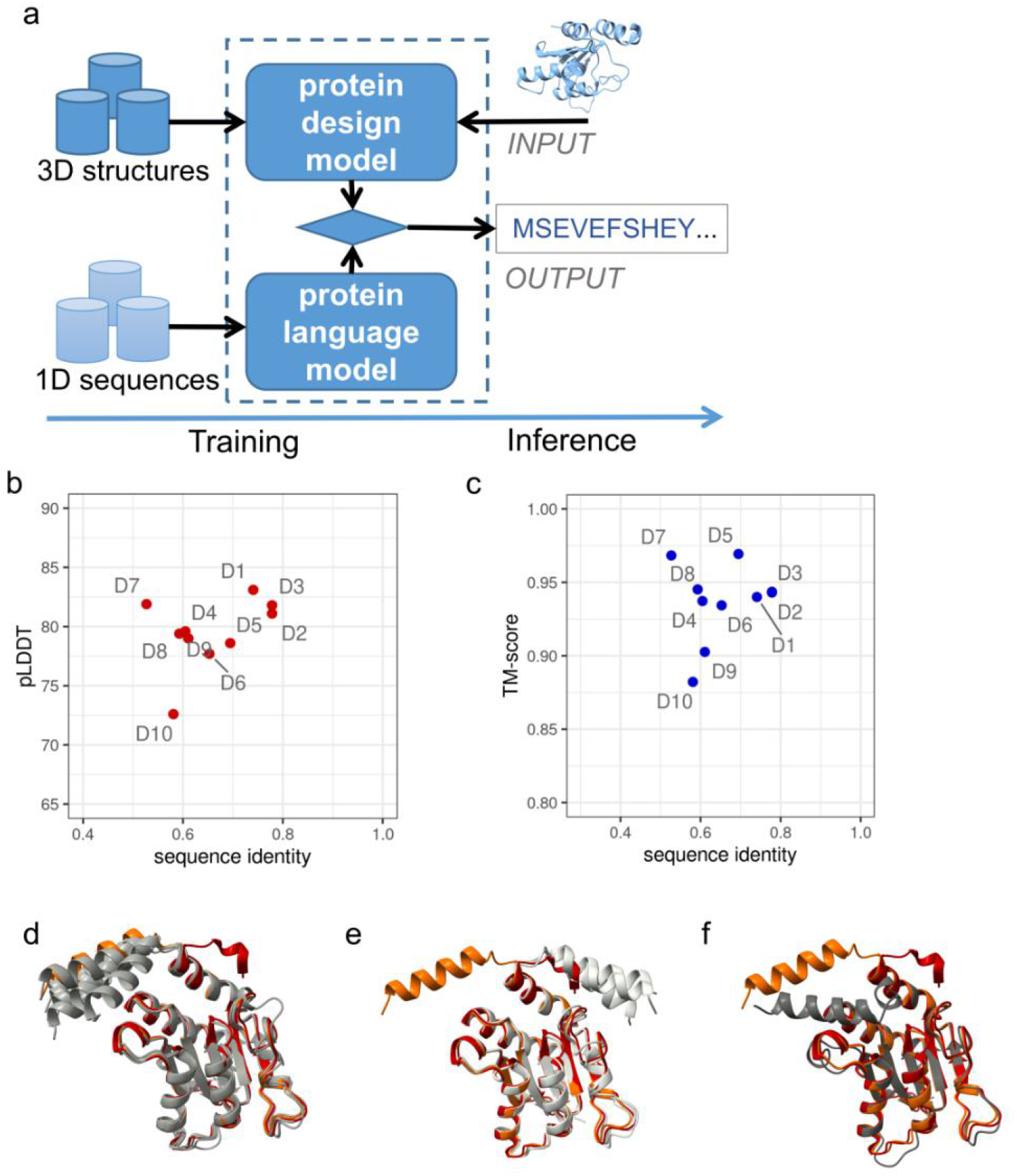
Protein design for adenine deaminase enzyme. (a) The schematic diagram of the protein design framework. (b) ESMFold was used to predict structures for 10 designed proteins (D1 to D10). The scatterplot shows the distribution of mean pLDDT and the sequence identity to TadA-8.20. (c) The scatterplot shows the distribution of TM-score and the sequence identity to TadA-8.20. (d) Predicted structures of designed proteins (D1-D4, D6 and D9, gray color) superimposed with TadA-8.20 real structure (red color) and predicted structure (orange color). (e) Predicted structures of designed proteins (D5, D7 and D8, gray color) superimposed with TadA-8.20 real structure (red color) and predicted structure (orange color). (f) Predicted structures of designed proteins (D10, gray color) superimposed with TadA-8.20 real structure (red color) and predicted structure (orange color).

To design adenine deaminase enzyme, we used TadA-8.20 structure^15^ (PDB code: 8e2p, chain A) as input. 10 designed sequences were selected after a strict filtering strategy. We used ESMFold^16^ predict the structures of these 10 designed sequences. All the structures were predicted confidently (mean pLDDT above 70, Figure 1b), and fold similarly with TadA-8.20, with the TM-score ranging from 0.88 to 0.97 (Figure 1c). The sequence identity between designed enzymes and TadA-8.20 are slow, ranging from 52.7% to 77.8%. The predicted structure of the TadA-8.20 fold similarly as the resolved structure except the C-terminal tail. According to the C-terminal differences, the 10 predicted structures could be classified to 3 classes. Class I including D1-D4, D6 and D9 are more like the predicted structure of the TadA-8.20 (Figure 1d). Class II including D5, D7 and D8 are more like the real structure of the TadA-8.20 (Figure 1e). D10 diverges from both the predicted and real structure of TadA-8.20 (Figure 1f).

### Experimental validation of the designed ABEs

We next test whether the designed enzymes have activity of gene base editing in human cells in the context of ABE. Human codon optimized sequences were cloned into ABE8.20-m plasmid, replacing the original tadA-8.20 sequence (Figure 2a). We co-transfected HEK293T cells with the ABE plasmid and single-guide RNA (sgRNA) expression plasmid targeting a previously characterized target site (Supplemental Table S1, site1). DNA editing efficiency was estimated using next generation sequencing three days after transfection.

**Figure 2.**
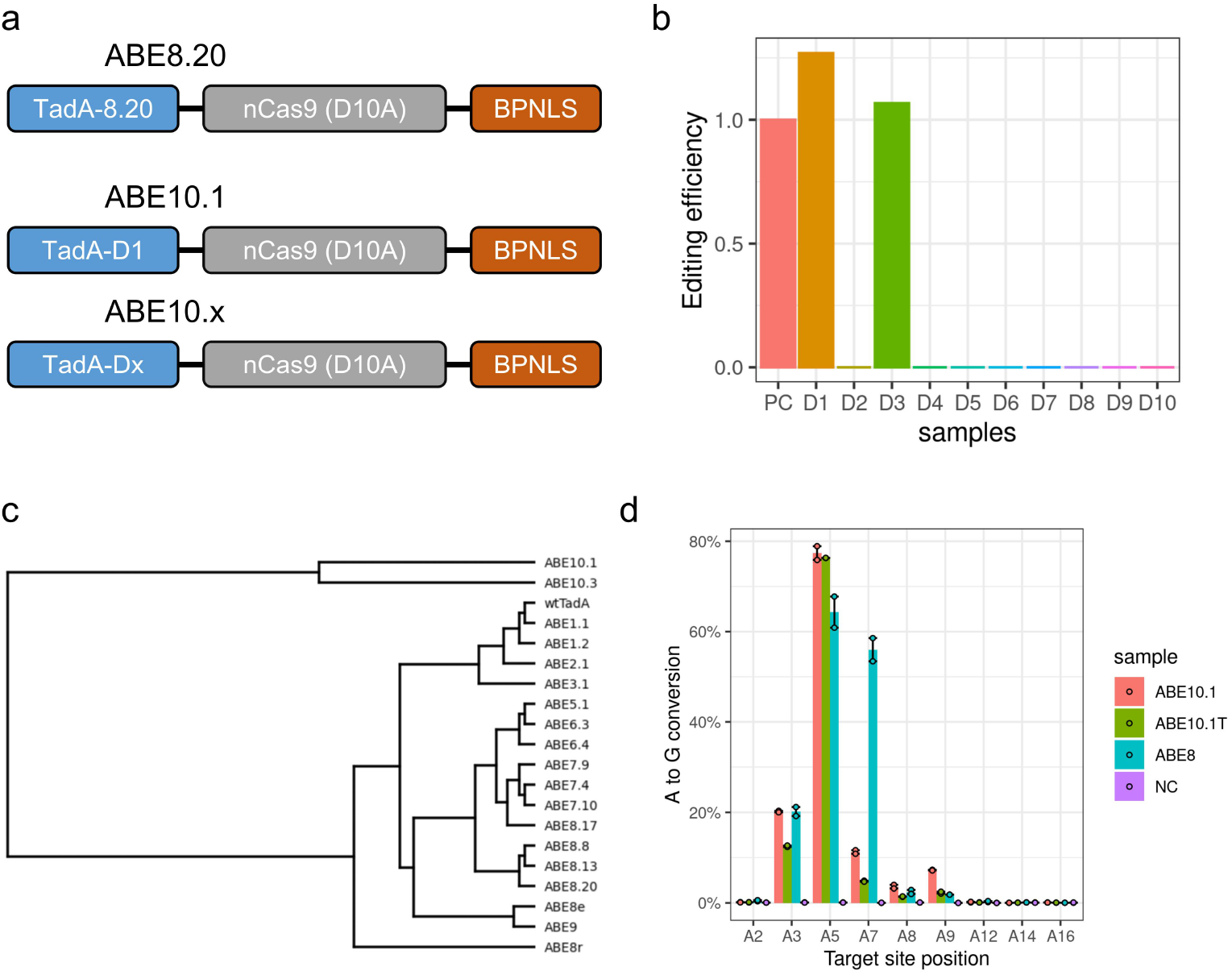
Experimental validation of the designed ABEs. (a) Architecture of ABE8.20 and ABE10s. The ‘x’ numerical value of ABE10.x indicates which mutations are included in the designed TadA of the corresponding editor. (b) Editing efficiency of the design ABEs at site1, normalized with the positive control (PC) of ABE8.20. (c) The phylogenetic tree of previous ABEs and the designed ABEs. Distances were calculated based on the protein sequences of TadAs. (d) Percentage of A to G conversion within the editing window of target site1.

Two out of the 10 variants have base editing activity. D1 and D3 shows 1.27 and 1.07 fold higher base editing efficiency at the test target site compared with ABE8.20, respectively (Figure 2b). Interestingly, these two designs belong to Class I of the designs. This observation indicates higher successful rate may be obtained with stricter structure alignment filters, especially when the predicted structures have systematic bias. Based on sequence alignment of TadAs (Supplemental Figure S1), we drew the phylogenetic tree of previous ABEs and the designed ABEs with base editing activity (Figure 2c). ABE10.1 and ABE10.3 show much larger divergence compared with the previous ABEs. This demonstrated that our protein design framework indeed found active enzymes in a much larger protein space.

Since ABE10.1 shows higher editing rate, we focus on this variant in the following study. To further characterize the effective editing window of ABE10.1, we check the editing rate at each A in target site. ABE10.1T (Figure S2), which contains a truncated version of D1, was also included in this analysis. Both ABE10.1 and ABE10.1T show statistically significant higher editing rate at the A5 peak locus, and statistically significant lower editing rate at the A7 locus. Both ABE10.1 and ABE10.1T, especially ABE10.1T has narrower editing window compared with ABE8.20 at this target site. This result also indicates the N- and C-terminal of TadA may have effect on the effective window size of ABEs.

### ABE10 shows improvement of base editing efficiency at multiple human genome sites

We further tested the ABE10.1 with 5 sgRNAs (Table S1, site2-site6) that are previously reported^15^. To further characterize the targeting abilities of adenine base editing, we treated HEK293T cells with ABE10.1, ABE10.1T or ABE8.20 that each target several endogenous genomics sites and measured A to G base conversion efficiency after three days (Figure 3a-3e). The increased editing efficiency of ABE10.1 editor compared with ABE10.1T was most evident when the editing levels within each protospacer were examined for all five sites. This indicates that the truncation of both N-ad C-terminal of TadA impairs the editing activity. The lower editing efficiency of ABE10s at A8 and A11 of site2 compared with ABE8.20 is consistent with our previous observation that ABE10s have narrower editing window, with peak at A5. For two out of the five sites, both ABE10.1 and ABE10.1T show increased editing efficiency at the most efficiently edited A within each protospacer. ABE10.1 and ABE10.1T achieve 66.7% and 56.6% editing efficiency at site4, which is 2.0 and 1.7 fold compared with ABE8.20, respectively (Figure 3d). For site5, ABE10.1 and ABE10.1T achieve 83.4% and 66.6% editing efficiency, which is 1.8 and 1.4 fold compared with ABE8.20, respectively (Figure 3e).

**Figure 3.**
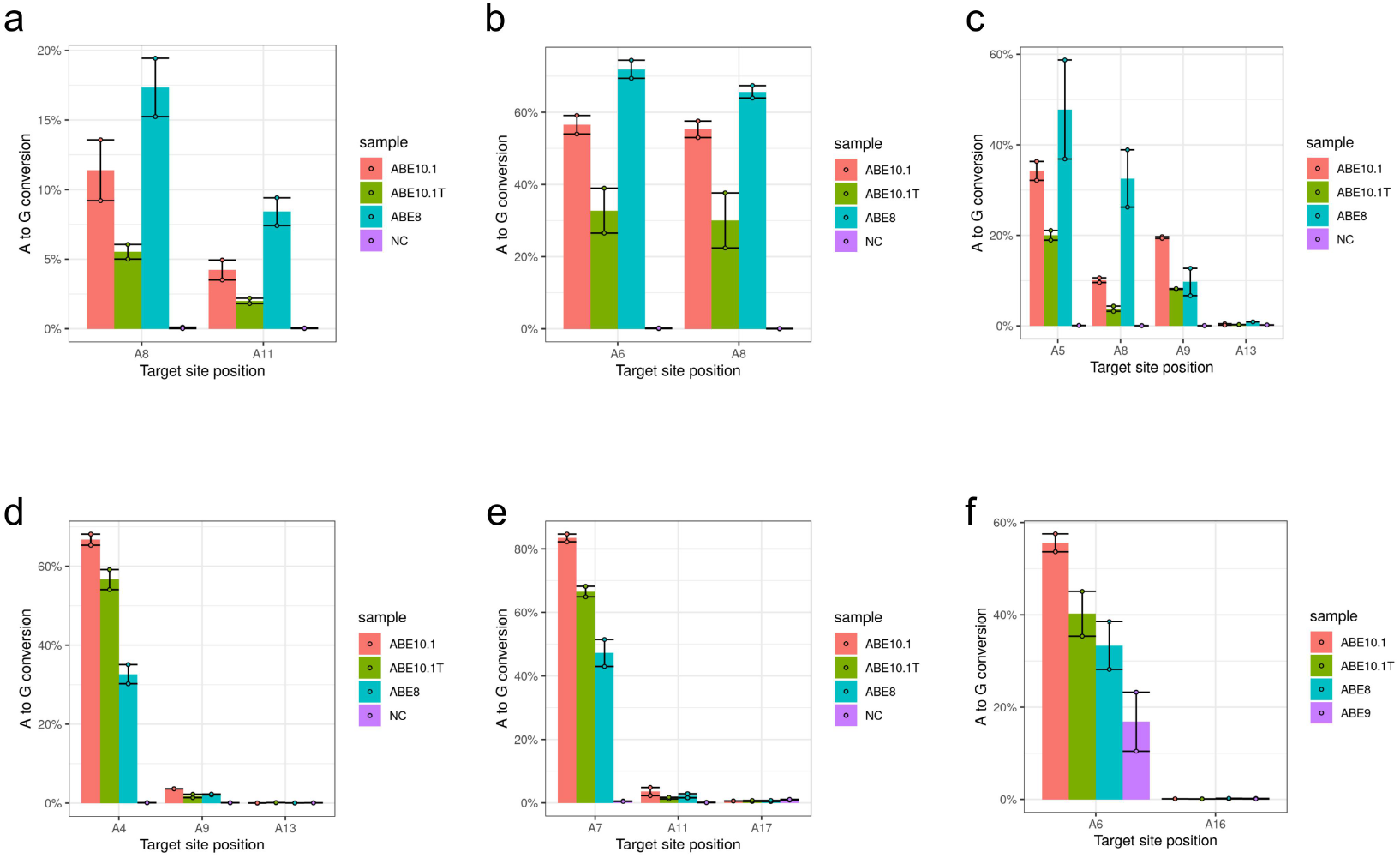
Base editing efficiency of ABE at multiple human endogenous genomics sites. (a) Editing efficiency of ABEs at site2. Wild type HEK293T cells were used as negative control (NC). (b) Editing efficiency at site3 (c) Editing efficiency at site4 (d) Editing efficiency at site5 (e) Editing efficiency at site6 (f) Editing efficiency at site7, which targets PCSK9

Since the designed ABE10s have not been optimized with any wet experiment, these results encourage us to ask whether ABE10s also have improved editing efficiency when targeting pathogenic mutations or therapeutic targets. The PCSK9 gene is a candidate target for in vivo gene editing. PCSK9 is a negative regulator of low-density lipoprotein (LDL), and rare gain-of-function mutations in human PCSK9 cause familial hypercholesterolaemia. We treated HEK293T cells with ABE10.1, ABE10.1T, ABE9 or ABE8.20 that each targets splice-donor site at the boundary of Pcsk9 exon 1 and intron 1 with a previously reported sgRNA^17,18^ (Table S1, site7). A to G base conversion efficiency was measured three days after transfection. ABE10.1 and ABE10.1T achieve 55.6% and 45.1% editing efficiency, which is 1.7 and 1.2 fold higher compared with ABE8.20, respectively (Figure 3f), whereas ABE9 has editing efficiency of 16.8%. This indicates ABE10s could be valuable alternative gene base editing tools.

### Bystander effect and off-target analysis of ABE10

Several studies have shown that some ABEs exhibit cytosine deamination activity which enables C-to-T/G/A conversions^8,19^. It is critical to reduce or eliminate both cytosine bystander effects and off-targeting editing of ABEs. To test the bystander effect, we test a previously reported target site^8^ (Table S1, site8). We first measured the base editing efficiency of ABE10.1, ABE10.1T or ABE8 at an endogenous genomic locus (FANCF gene, site8) for both adenine and cytosine editing in HEK293T cells. While ABE10.1 and ABE10.1T achieve 48.9% and 44.9% editing efficiency respectively, ABE8.20 has 34.9% editing efficiency at the desired A4 of target site (Figure 4a). The cytosine bystander editing at C6 of target site for ABE10.1, ABE10.1T and ABE8.20 is 3.6%, 2.5% and 18.0%, respectively. ABE10.1 and ABE10.1T strikingly decreased the cytosine bystander mutation rate by 5.0-fold and 7.2-fold in comparison to ABE8.20.

**Figure 4.**
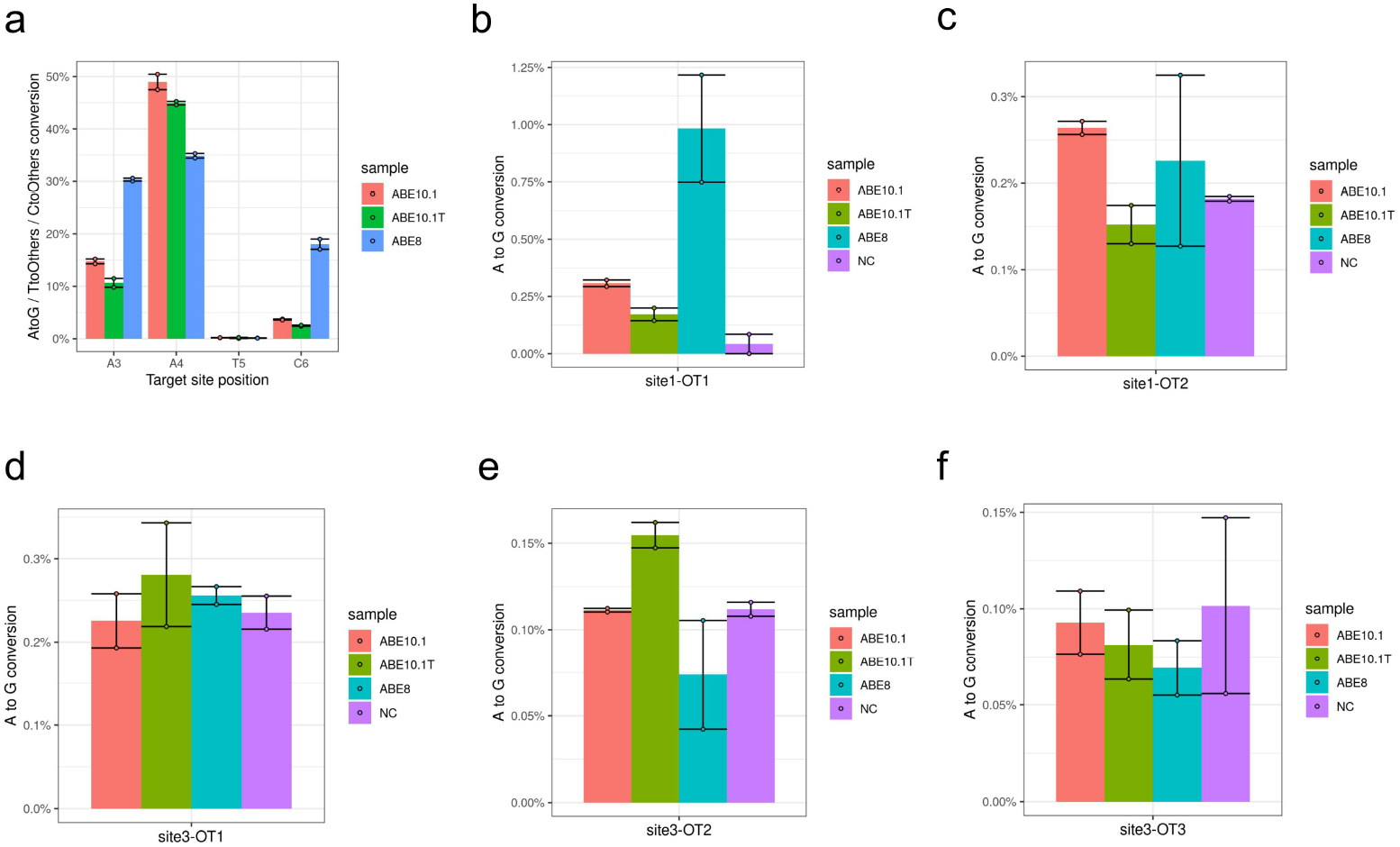
Bystander effect and off-target analysis. (a) Editing efficiency of ABEs at site8 which targets FANCF gene. (b) Off-target editing efficiency at site1-OT1, Wild type HEK293T cells were used as negative control (NC) (c) Off-target editing efficiency at site1-OT2 (d) Off-target editing efficiency at site3-OT1 (e) Off-target editing efficiency at site3-OT2 (f) Off-target editing efficiency at site3-OT3

To characterize off-target editing of ABE10, we analyzed gRNA-dependent off-target base editing frequency of ABE10 in HEK293T cells. gRNAs for which the gRNA off-target profile has been previously reported with Cas9 and base editors were used^15^. We tested the top 2 known off-target sites for site1 and the top 3 off-target sites for site3. In general, all three ABEs tested have very low off-target (OT) editing for these sites. For OT1 of site1 (site1-OT1), ABE10.1 and ABE10.1T have lower editing rate than ABE8.20 (Figure 4b). For other 4 off-target sites, the editing rate is comparable to the background level of random mutations (Figure 4c-4f). Although we have not detected significant off-target editing in these sites. further off-target analysis will be needed in the near future, especially for clinical applications.

## Discussion

Gene base editing technologies derived from directed evolution of natural systems and mutant-testing strategies have enabled programmable and precise manipulation of the genome. Though powerful, these strategies explore the local protein space to find an improved solution to specific objective function. Applications of AI technologies such as protein design models and protein language models in biological systems provide an effective strategy to explore a much larger protein space that were never touched by natural evolution and directed evolution.

In this work, we designed a novel adenine deaminase enzyme with our protein design framework, which integrated both protein sequence design model and protein language model, and created new adenine base editors, ABE10s. Instead of introducing the technical details of our protein design framework, we focused on characterizing the novel AI-designed adenine deaminase enzyme and the derived ABE10s. Surprisingly, without any wet experimental optimization, the AI-designed ABE10s has promising performance. The ABE10s show high editing efficiency with reduced bystander editing and off-target editing rate. At five out of the eight target sites tested, ABE10.1 show even much higher on-target editing efficiency compared to the well-optimized ABE8.20. To our knowledge, this is the first AI-derived adenine deaminase enzyme and ABE that shows higher editing efficiency compared to the state-of-the-art ABE8.20.

Further optimization of ABE10s with directed evolution, rational design and/or high-throughput screening has high potential to discover improved variants. Alternatively, sampling other protein space with our framework could also lead to discovery of other diverse enzymes with different characteristics. ABE10 includes two key components: the editing component, which is the AI-derived TadA and the guiding component, the widely used SpCas9n/sgRNA pair. A promising direction is to design a novel guide system, which is not limited to Cas or TnpB proteins. The ideal guide system will be safe, precise, effective and compact (SPEC. for short). This poses new challenges for the design of complex system with multiple tasks to protein design community.

## Author contributions

G.L. and Y.Y. conceived of the ideas implemented in this project. G.L. and Y.Y. designed the models and sequences. Y.C. and R.L. performed the wet experiments, Q.G. and Y.N.Y. assisted with wet experiments. G.L. analyzed the data and results, G.L. supervised the project. G.L., Y.C. and Y.Y. wrote the paper.

## Methods

### Protein design framework

The protein design framework contains three key parts: a protein design model and a protein language model are integrated with a fusion module. The protein design model was trained as a fixed-backbone protein sequence design task, where the inputs are protein structures and outputs are protein amino acid sequences expected to fold into the input structures. The protein language model was trained as a masked language model task, which predicts the amino acid types of the random masked tokens in the input sequences. A fusion module that takes the output head of two models as input and predict the distribution of amino acid types for the protein of interest.

### Structure and sequence analysis

3D structures were predicted with ESMFold and the pLDDT scores were reported by ESMFold. 3D structures visualization and structure alignment were supported by ChimeraX^25^. Sequence identity and the phylogenetic tree was calculated using biotite package. TM-score was calculated with TM-align^20^.

### Next generation sequence analysis

Editing frequency was estimated using our bioinformatics pipline. Briefly, trimmomatic^21^ was used to to cut adapters to get clean reads. Bowtie2^22^ was used to align reads to hg38 reference genome. Samtools^23^ was used to transform the bam/sam format. Bam-readcount^24^ was used to calculate the read depth and mutation type and mutation frequency at each bases within the target sites.

### Cell culture

HEK293T cell were obtained from the cell bank(CAS) and cultured in 90% DMEM(Gibco), supplemented with 10% Fetal Bovine Serum(Gibco) and 1% penicillin/streptomycin (Invitrogen). These cell were grown at 37 °C and 5% CO2 and routinely passaged using TrypLE™ Express (Gibco).

### Plasmid Construction of new base editors

First, the designed new tadA sequences were synthesized commercially through the regular gene synthesis of Azenta life science. We used the Phanta Max Super-Fidelity DNA Polymerase kit(Vazyme) to amplify the pre-designed fragments from the gene synthesis universal vector pUC-GW-Amp and add the homology arm for later homologous recombination. The sequences of the homology arms were forward 5’-3’ atacgactcactatagggagagccgccacc, reverse 5’-3’ tgccgctagaaccaccagaagaaccaccaga. The 500bp PCR fragments were purified by DNA electrophoresis and gel extraction(TaKaRa MiniBEST Agarose Gel DNA Extraction Kit Ver.4.0). The insert fragments for homologous recombination were obtained.

Then, linearized editing vectors was linearized from ABE8.20m vector(addgene) by reverse PCR. The target fragments were cut off after agarose gel electrophoresis. Gel extraction and purification were performed to obtain the vector fragments for homologous recombination.

At last, we used NEBbuilder HiFi DNA Assembly Master Mix to perform homologous recombination integration of the above insert fragments and vector fragments, with a molar ratio of vector fragments to insert fragments of 1:10. Then we transformed the integration products into the competent cell trans-5α(TransGen biotech). The next day, 5-10 monoclonal strains were picked and identified by Sanger sequencing. Finally, we picked the clones that matched the pre-designed sequences exactly for amplification and extracted the plasmid DNA for subsequent single-base editing experiments.

### Cell transfection

HEK293T cells were seeded initially in a 24-well plate to ensure 50%-60% confluence on the second day of the experiment. After 12h to 16h hours, lipo2000 reagent was applied for transfection. For a 24-well plate, the amount of transfection reagent is as follows Tube A: optiMEM 25ul+lipofectamine2000 3ul, Tube B: opti-MEM 25ul+ 2ug DNA (1.5ug ABE +0.5ug sgRNA). Then, we gently mixed tubes A and B, let stand for 15 minutes, and evenly spot on a 24-well plate, incubate for 72h-120h. After transfection, Cell were collected for measuring base editing efficiency.

### Genome extraction, target sequence amplification and sequencing

Genomic DNA was extracted by Qiagen tissue&blood DNA extraction kit. Then two-step PCR methods were used to construct next generation sequencing libraries. The first step amplified the target fragments and the second step added Tsingke FastNGS adapter(5’ end TCGTCGGCAGCGTCAGATGTGTATAAGAGACAG, 3’ end GTCTCGTGGGCTCGGAGATGTGTGTATAAGAGACAG) using Vazyme Phanta Max Super-Fidelity DNA Polymerase kit. Finally, the gel extracted and purified products were sent to FastNGS (Tsingke Biotech) for high-throughput sequencing with a sequencing depth of 5000x.

