## supplemental for "An AI-designed adenine base editor"

### Supplementary information

#### Table S1. sgRNA sequences

| | | | |
|---|---|---|---|
| 1 | Site1-sense | GAACACAAAGCATAGACTGC | |
| 2 | Site2-sense | GAGTCCGAGCAGAAGAAGAA | EMX1 |
| 3 | Site3-sense | GGCCCAGACTGAGCACGTGA | |
| 4 | Site4-sense | GCCCAGCAATTCACTGTGAAG | |
| 5 | Site5-sense | GTCATCTTAGTCATTACCTG | RNF2 |
| 6 | Site6-sense | GGTCGTAGCCAGTCCGAACCC | EMX1 |
| 7 | Site7-sense | CCCGCACCTTGGCGCAGCGG | PCSK9 |
| 8 | Site8-sense | GGAATCCCTTCTGCAGCACC | FANCF |

#### Table S2. PCR primer sequences for on-target sites

| | | |
|---|---|---|
| 1 | Site1-for | CCAGCCCCATCTGTCAAACT |
| 2 | Site1-rev | TGAATGGATTCCTTGGAAACAATGA |
| 3 | Site2-for | CAGCTCAGCCTGAGTGTTGA |
| 4 | Site2-rev | CTCGTGGGTTTGTGGTTGC |
| 5 | Site3-for | ATGTGGGCTGCCTAGAAAGG |
| 6 | Site3-rev | CCCAGCCAAACTTGTCAACC |
| 7 | Site4-for | AGGTGGGGGTGACTCCTTTTTTGGA |
| 8 | Site4-rev | GGGCAGAAGGAAAAATCTATCCTGGAA |
| 9 | Site5-for | ACGTCTCATATGCCCCTTGG |
| 10 | Site5-rev | ACGTAGGAATTTTGGTGGGACA |
| 11 | Site6-for | GCTGCTGGAATACCGAGGAC |
| 12 | Site6-rev | GCAACTCTCTTTTCTCCGGGA |
| 13 | Site7-for | AGGACGAGGACGGCGACTAC |
| 14 | Site7-rev | AAACTGAGGCCCGAGAGG |
| 15 | Site8-for | AAGGAACACGGATAAAGACGCTGGG |
| 16 | Site8-rev | TAGGTAGTGCTTGAGACCGCCAGAA |

#### Table S3. PCR primer sequences for off-target sites

| | | |
|---|---|---|
| 1 | Site1-OT1-for | GTGTGGAGAGTGAGTAAGCCA |
| 2 | Site1-OT1-rev | ACGGTAGGATGATTTCAGGCA |
| 3 | Site1-OT2-for | CACAAAGCAGTGTAGCTCAGG |
| 4 | Site1-OT2-rev | TTTTTGGTACTCGAGTGTTATTCAG |
| 5 | Site3-OT1-for | TCCCCTGTTGACCTGGAGAA |
| 6 | Site3-OT1-rev | CACTGTACTTGCCCTGACCA |
| 7 | Site3-OT2-for | TTGGTGTTGACAGGGAGCAA |
| 8 | Site3-OT2-rev | CTGAGATGTGGGCAGAAGGG |

| | | |
|---|---|---|
| 9 | Site3-OT3-for | TGAGAGGGAACAGAAGGGCT |
| 10 | Site3-OT3-rev | GTCCAAAGGCCCAAGAACCT |

#### Figure S1. Sequence alignment of TadAs

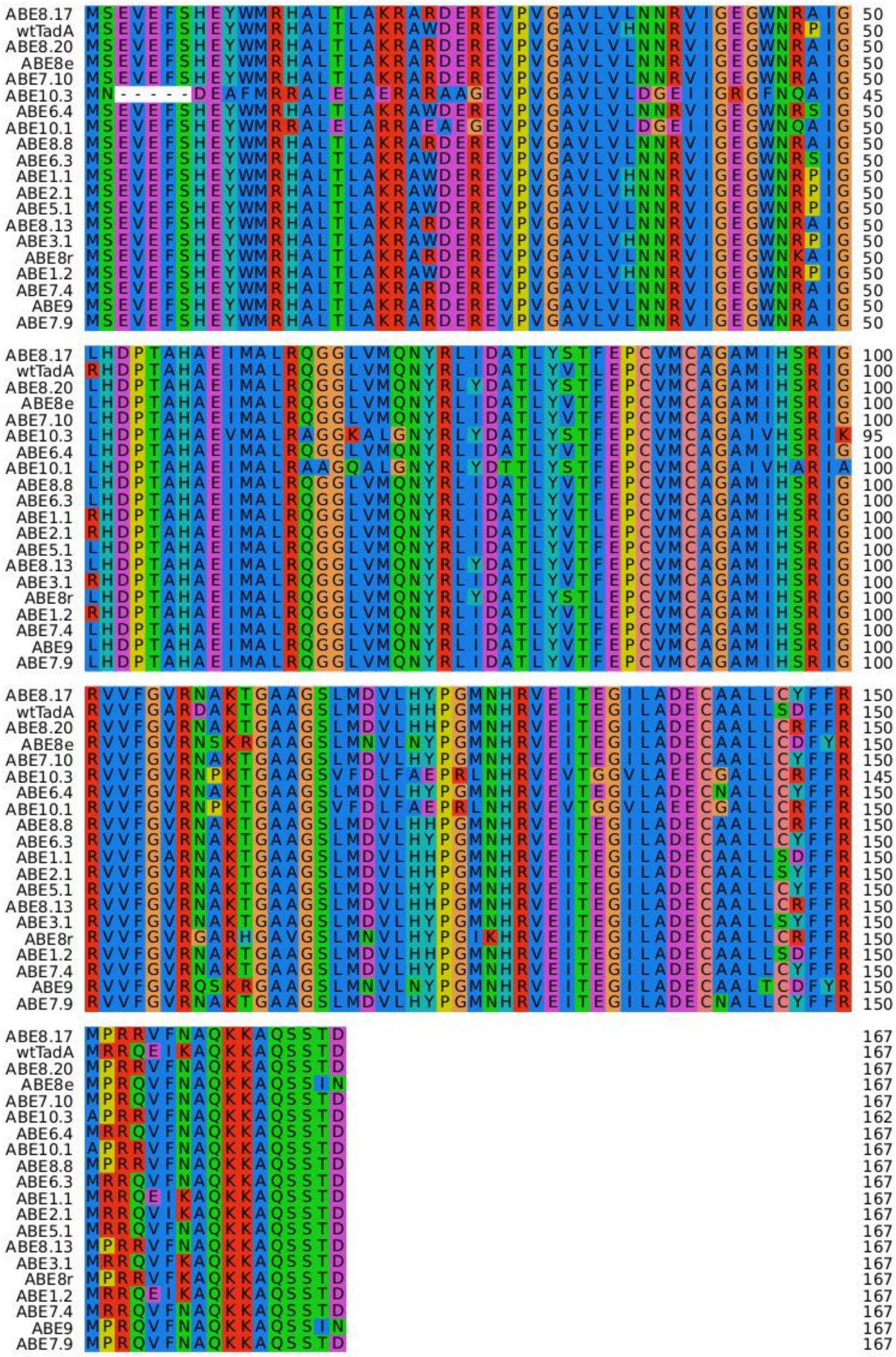

**Figure S2. Sequence alignment between TadA of ABE10.1 and ABE10.1T**

```
ABE10.1   MSEVEFSHEYWMRRALELARRAEAEGEVPVGAVLVLDGEIIGEGWNQAIG  50
ABE10.1T  -----------MRRALELARRAEAEGEVPVGAVLVLDGEIIGEGWNQAIG  39

ABE10.1   LHDPTAHAEIMALRAAGQALGNYRLYDTTLYSTFEPCVMCAGAIVHARIA  100
ABE10.1T  LHDPTAHAEIMALRAAGQALGNYRLYDTTLYSTFEPCVMCAGAIVHARIA  89

ABE10.1   RVVFGVRNPKTGAAGSVFDLFAEPRLNHRVEVTGGVLAEECGALLCRFFR  150
ABE10.1T  RVVFGVRNPKTGAAGSVFDLFAEPRLNHRVEVTGGVLAEECGALLCRFFR  139

ABE10.1   APRRVFNAQKKAQSSTD                                  167
ABE10.1T  APRRVFN----------                                  146
```
